# Age-related differences in the association between REM sleep and the polygenic risk for Parkinson’s disease

**DOI:** 10.1101/2025.02.05.636657

**Authors:** Puneet Talwar, N. Mortazavi, Ekaterina Koshmanova, Vincenzo Muto, Aurora Gasparello, Christian Degueldre, Christian Berthomier, Fabienne Collette, Christine Bastin, Christophe Phillips, Pierre Maquet, Zubkov Mikhail, Gilles Vandewalle

## Abstract

**Objective:** Parkinson’s disease (PD) is one of the rare diseases for which sleep alteration is a true marker of disease outcome. Yet, how the association between sleep and PD emerges over the healthy lifetime is not established. We examined association between the polygenic risk score (PRS) for PD and the variability in the electrophysiology of Rapid Eye Movement (REM) sleep in 433 younger (18-31y) and 85 late-midlife (50-69y) healthy individuals.

**Methods:** In this prospective cross-sectional study, in-lab EEG recordings of sleep were recorded to extract REM sleep metrics. PRS was computed using SBayesR approach.

**Results:** Generalized Additive Model for Location, Scale and Shape (GAMLSS) analysis showed significant association of REM duration (**p_corr_=0.03)** and theta energy in REM (**p_corr_=0.004**) with PRS for PD in interaction with age group. In the younger sub-sample, REM duration and theta energy were positively associated with PD PRS. In contrast, in the late-midlife sub-sample, the same associations were negative (though only qualitatively for REM theta energy) and may differ between men and women.

**Interpretation:** REM sleep is associated with the PRS for PD in early adulthood, 2 to 5 decades prior to typical symptoms onset. The association changes from positive in younger individuals, presumably free of alpha-synuclein, to negative in late-midlife individuals, possibly because of the progressive presence of alpha-synuclein aggregates or of the repeated increased oxidative metabolism imposed by REM sleep. Our findings may unravel core associations between PD and sleep and may contribute to novel intervention targets to prevent or delay PD.

---

Parkinson’s disease (PD) is the second most prevalent neurodegenerative disorder. The global burden of PD has surged from 2.5 million people in the 1990s to 8.5 million today, a trend expected to continue in the next decades^1^. The incidence of PD increases with age and is higher in men than women^2^. PD is characterized by a heterogeneous pathology, with significant variations in clinical manifestations - with both motor and non-motor symptoms and signs, including during sleep - as well as in disease progression, and responses to treatment^2, 3^. The neuropathological hallmark of PD consists of conspicuous lesions in the substantia nigra (SN), a major dopaminergic nucleus forming the nigrostriatal pathway^4^, in the form of intracellular inclusions of α-synuclein known as Lewy bodies, accumulation of neuromelanin and iron and neuronal loss.

Currently, PD remains a clinical diagnosis, as no laboratory or neuroimaging biomarkers can definitively confirm the disease. Existing pharmacological treatments primarily address motor symptoms without halting the underlying neurodegenerative processes^5^. The challenge in developing neuroprotective therapies is compounded by the extensive neurodegeneration present by the time motor symptoms and PD diagnosis occur. Therefore, early identification of presymptomatic PD, or of high risk for PD, is a critical medical need to develop effective neuroprotective strategies, delay or prevent the symptomatic stage of the disease.

Excessive daytime sleepiness significantly increases the risk of developing PD^6^. Critically, the majority of individuals with idiopathic REM sleep behavior disorder (RBD)— characterized by loss of normal atonia and vigorous movements during REM sleep—progress to PD or cognitive impairment within 10-15 years, making RBD a prominent sleep-related risk factor for PD^6^. The progressive disruption of REM/NREM sleep transitions in idiopathic RBD and PD likely reflects the early involvement of subcortical regions, such as the locus coeruleus (LC)^6–8^. The literature highlights a bidirectional detrimental relationship between sleep alterations and neurodegeneration, including in PD^9^. The recently identified glymphatic system, which is proposed to be active during sleep and suppressed by LC noradrenaline, appears crucial for α-synuclein clearance as its dysfunction seem to exacerbate α-synuclein accumulation^6, 10^. Sleep alterations, in REM sleep, may therefore not only provide early means to assess the risk for developing PD but may also provide novel intervention targets (as one can act on sleep). However, which key features of sleep should be monitored is not established.

Although monogenic forms account for 3-5% of PD cases, recent genome-wide association studies (GWAS) indicate that idiopathic sporadic PD is highly polygenic^11, 12^. Ninety genetic risk variants collectively account for 16-36% of the heritable risk of non-monogenic PD^3^. Consistent with the common disease-common variant hypothesis, PD genetic risk results from the synergistic effect of numerous common low-risk variants – in addition to environmental influences. Polygenic risk scores (PRS), derived from GWAS, hold promise for predicting and stratifying PD risk in asymptomatic individuals, and already provide a critical tool to investigate the neurobiology of the disease in individuals of any age, i.e. prior to PD hallmarks can be detected and free from disease co-morbidity^12^. In recent years, polygenic risk score for PD has been associated with olfactory impairment^13^, retinal structural integrity^14^, cognitive deficits^15^. In this cross-sectional study, we aimed to investigate the relationship between Polygenic Risk Scores (PRS) for PD and REM sleep metrics in healthy young and late-midlife individuals. We analysed key REM sleep characteristics related to PD pathophysiology, including metrics associated with REM sleep duration, intensity and continuity^16^. Although the literature on the associations between sleep metrics and PD PRS are scarce, based on the age-related changes in brain integrity we anticipated that associations would be different in the two age groups.

## Results

In the present study, we collected the habitual night time sleep of 518 healthy individuals under EEG (**Table 1**), in two sub-sample of younger -18 to 31y; N = 433; 44 women - and late-midlife -50 to 69y; N=85; 58 women-collected from seven previous multi-modal cross-sectional projects^17–19^. We quantified four REM sleep metrics based on their potential association with PD: 1) REM duration; 2) REM latency; 3) number of arousals during REM sleep, to characterise sleep continuity^20^; 4) REM theta energy, i.e. the cumulated theta power during REM sleep, associated with REMS intensity over its most typical oscillatory activity^20^. By *a priori* selecting variables of interest, we reduced the multiple comparison burden. We further used the summary statistics of one of the largest PD-GWAS available to date (N = 482,730)^21^ to compute individual PRS for PD in our sample – based on DNA extracted from blood samples - and related these to sleep EEG characteristics. The overview of the study design is provided in **Fig. 1**.

**Figure 1:**
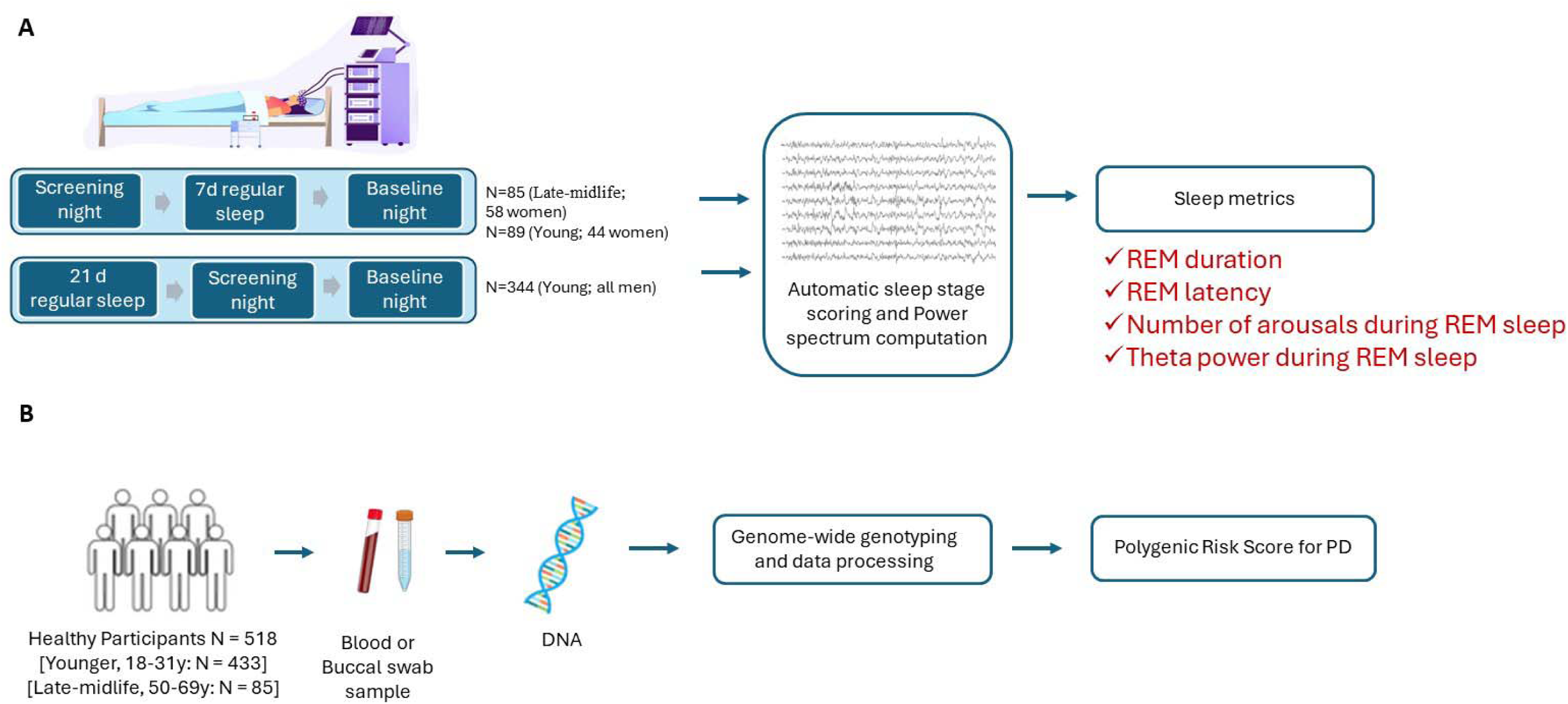
Overview of the study design: **A.** In-lab recordings of habitual sleep to extract REM sleep macro and microstructure metrics. **B.** Parkinson’s disease polygenic risk score (PRS) computation using available GWAS data and DNA extracted from blood sample and PD summary statistics.

**Table 1.** Characteristics for the total sample as well as young and late-midlife subgroups.

| Characteristic | All | Young<br>Mean (SD) | Late-midlife<br>Mean (SD) | p-value<sup>1</sup> |
| --- | --- | --- | --- | --- |
| Sample size (N) | 518 | 433 | 85 |  |
| Sex (Men) | 80.31% | 89.84% | 31.76% | <0.001 |
| Age (years) | 28.17 (14.09) | 22.10 (2.69) | 59.09 (5.26) | <0.001 |
| BMI (kg*m<sup>-2</sup>) | 22.55 (2.61) | 22.11 (2.32) | 24.82 (2.85) | <0.001 |
| Anxiety<sup>b</sup> | 2.67 (3.10) | 2.64 (3.16) | 2.78 (2.86) | 0.7 |
| Mood<sup>b</sup> | 3.29 (3.77) | 2.91 (3.41) | 4.85 (4.66) | <0.001 |
| Sleep Quality<sup>a</sup> | 3.71 (2.08) | 3.41 (1.72) | 4.89 (2.86) | <0.001 |
| Daytime sleepiness<sup>a</sup> | 5.95 (3.62) | 5.91 (3.53) | 6.13 (3.98) | 0.6 |
| Chronotype<sup>a</sup> | 50.87 (8.35) | 50.21 (8.35) | 53.49 (7.87) | <0.001 |
| Total Sleep Time (min) | 439.30 (47.11) | 448.65 (42.39) | 391.65 (40.88) | <0.001 |
| Sleep onset latency (min) | 15.87 (11.58) | 16.41 (11.68) | 13.12 (10.74) | 0.015 |
| REM duration (min) | 114.89 (28.55) | 119.14 (26.46) | 93.23 (29.17) | <0.001 |
| REM percentage (%) | 26.01 (5.54) | 26.47 (5.10) | 23.65 (6.93) | 0.001 |
| REM arousal (N) | 45.83 (24.18) | 51.12 (22.36) | 18.89 (12.13) | <0.001 |
| REM latency (min) | 92.04 (42.89) | 93.65 (41.38) | 83.85 (49.33) | <0.001 |
| REM delta power (0.5-4 Hz) (μV<sup>2</sup>) | 66,706<br>(69,597) | 72,453<br>(72,690) | 37,428<br>(40,005) | <0.001 |
| REM theta power (4-8Hz) (μV<sup>2</sup>) | 7,305 (5,273) | 7,885 (5,425) | 4,354 (3,039) | <0.001 |
| REM alpha power (8.25-12Hz) (μV<sup>2</sup>) | 4,192 (3,309) | 4,455 (3,426) | 2,850 (2,202) | <0.001 |
| REM sigma power (12.25-16 Hz) (μV<sup>2</sup>) | 1,287 (885) | 1,376 (919) | 832 (476) | <0.001 |
| NREM Slow wave energy (μV<sup>2</sup>) | 164,620<br>(163,436) | 180,901<br>(171,319) | 81,686<br>(71,642) | <0.001 |

<sup>1</sup>Pearson's Chi-squared test; Welch Two Sample t-test between young and late-midlife group Sleep quality was assessed by the Pittsburgh Sleep Quality index (PSQI). Daytime sleepiness was measured by the Epworth Sleepiness Scale, Chronotype was assessed by the Morningness-Eveningness Questionnaire (MEQ, ). Anxiety was estimated by the Beck Anxiety Inventory and Mood was estimated by the 21-item Beck Depression Inventory II; Total Sleep Time (TST) was extracted from polysomnography recordings. The data for Anxiety (N=428), Mood (N=428), Sleep quality (N=425), Daytime sleepiness (N=425) and Chronotype (N=425) excludes some young participant sub-sample due to unavailability.

### Associations between REM sleep metrics and age

We first assessed the differences between age groups and REM sleep metrics of interest through boxplots and t-test (**Fig. 2 and Table 1**). All four sleep metrics were significantly lower in the late midlife subgroup as compared to the young subgroup (p<0.001), which was expected based on previous literature, except for the while the number of REM arousals which was previously reported to increase significant^22, 23^.

**Figure 2:**
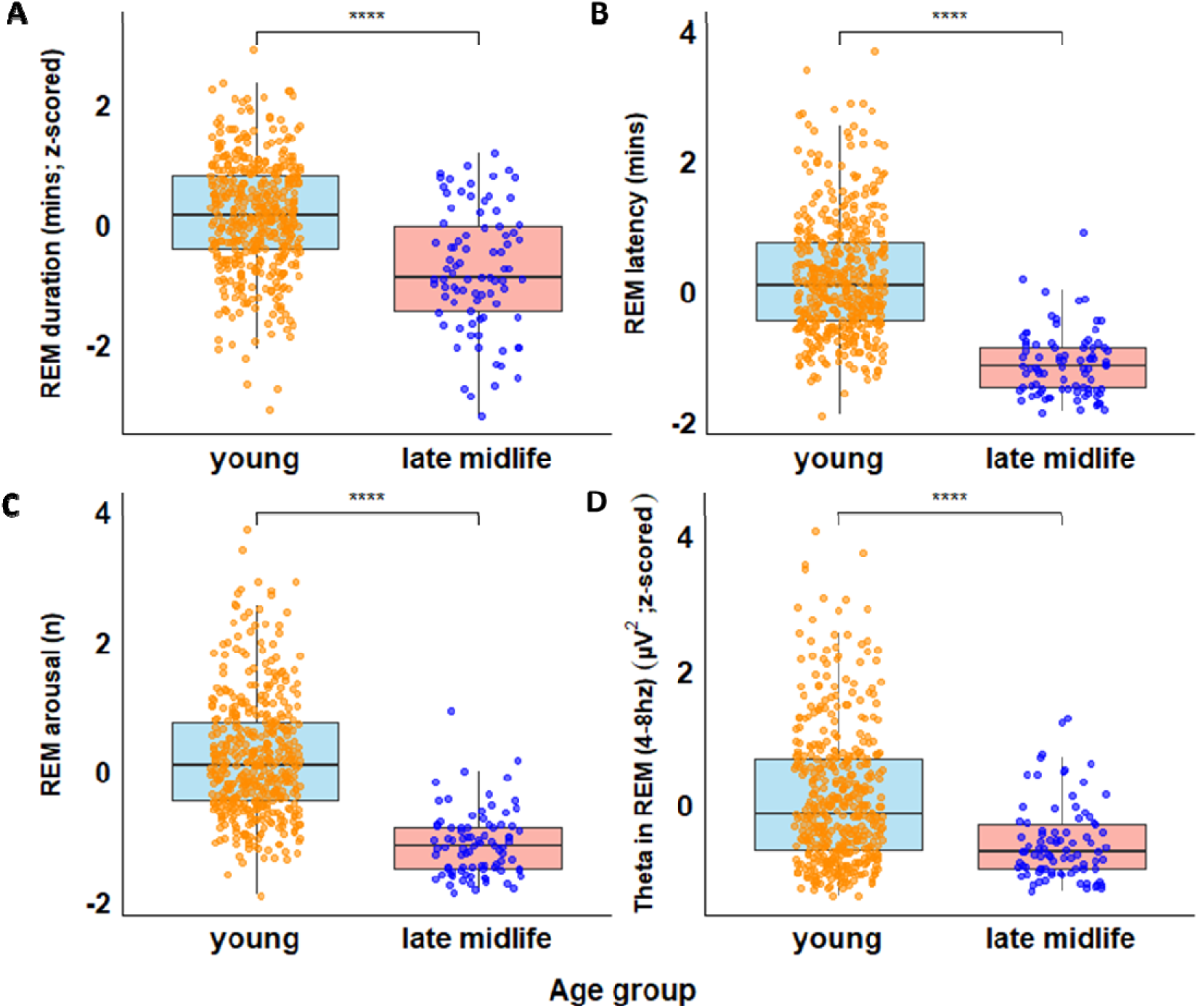
The boxplots for four sleep metrics (A-D) during baseline night and age groups (young and late midlife) with t-tests are reported. The sleep metrics were significantly higher in younger subgroup as compared to the late midlife group. *, **, *** refers to p values < .05, .01, .001.

### Associations between polygenic risk for PD with REM sleep duration and intensity

Our statistical analyses consisted of a generalized additive model for location, scale and shape (GAMLSS) which is a flexible distributional regression approach and is considered as an improvement and extension to the generalized linear models (GLM)^24^. Our primary GAMLSS regression analyses included each of the four sleep metrics of interest and PRS for PD with age group included as interaction variable with PRS. The GAMLSS with REM duration (p=.016, **p_corrected_=0.03)** and theta energy in REM (p=.001, **p_corrected_=0.004**) as dependent variables yielded significant interaction between PD PRS and age group after controlling for sex, body mass index (BMI) and total sleep time (TST) or REM duration (**Table 2**; – **Suppl. Fig S1** for display). REM latency and arousal in REM did not, however, reveal significant associations with PRS for PD (**Table 2**).

**Table 2.** Results derived from regression analysis when testing for associations between sleep parameters and PRS values computed for Parkinson’s disease risk with age group as an interacting variable (N=518).

| Sleep parameters | REMS duration |  | REMS theta Power |  | REMS latency |  | REMS Arousal |  |
| --- | --- | --- | --- | --- | --- | --- | --- | --- |
|  | Estimate<br>[95% CI] | p-value | Estimate<br>[95% CI] | p-value | Estimate<br>[95% CI] | p-value | Estimate<br>[95% CI] | p-value |
| PRS*Age group | 0.01<br>[0.00, 0.02] | **.016** | 0.04<br>[0.02, 0.06] | **.001** | -0.31<br>[-1.16, 0.54] | .477 | 0.01<br>[-0.00, 0.03] | .086 |
| PRS* | -0.01<br>[-0.01, 0.00] | .075 | -0.03<br>[-0.05, -0.01] | **.014** | 0.15<br>[-0.67, 0.96] | .725 | -0.01<br>[-0.003, 0.00] | .076 |
| Age group | -0.01<br>[-0.08, 0.06] | .847 | 0.41<br>[0.17, 0.64] | **.001** | 4.98<br>[-2.93, 12.90] | .216 | 0.62<br>[0.48, 0.76] | **<.001** |
| Sex | 0.01<br>[-0.04, 0.06] | .665 | -0.08<br>[-0.24, 0.09] | .346 | 3.59<br>[-2.75, 9.94] | .266 | 0.05<br>[-0.05, 0.16] | .302 |
| BMI | 0.003<br>[-0.00, 0.01] | .434 | 0.02<br>[-0.01, 0.04] | .165 | 0.29<br>[-0.66, 1.23] | .552 | -0.01<br>[-0.02, 0.00] | .176 |
| TST | 0.003<br>[0.00, 0.00] | **< .001** | 0.001<br>[0.00, 0.00] | **.030** | 0.05<br>[-0.00, 0.10] | .052 | - | - |
| REM duration | - | - | - | - | - | - | 0.01<br>[0.01, 0.01] | **< .001** |

PRS: polygenic risk score; BMI: body mass index; REMS: rapid eye movement sleep
Regression coefficients (log scale) with 95% confidence intervals are shown.
\*PRS was computed for SNP reaching p-value threshold of $1 \times 10^{-8}$ to restrict the number of SNPs included in the PRS estimation.

To gain insight into the interactions, we split our sample into the young (N=433) and late-midlife (N=85) sub-samples and recomputed post hoc GAMLSS in each sub-sample (see **Table 1** for demographic and sleep metrics in each subgroup). We found that while the association was significantly positive in the younger subsample for REM duration (**p=0.014**), i.e. higher REM duration is related to higher PD PRS, the association was significantly negative in the late-midlife subsamples (**p=0.02)** (**Table 3**, **Fig. 3A**). In the young sub-sample, GAMLSS yielded no sex specific association for REM duration with PD PRS (Suppl. Table S1 and Fig. S2A). However, when sex was included in the GAMLSS model including the late-midlife sub-sample, it yielded significant sex-by-PRS interaction (**p=0.03**) with negative and positive links, respectively, in men and women (**Suppl. Table S1**; significant association only with men for REM duration (p=0.005) (**Suppl. Table S2, Fig. 4A&C**). These findings suggest that REM duration is associated with PD PRS in an age and possibly sex dependent manner.

**Figure 3:**
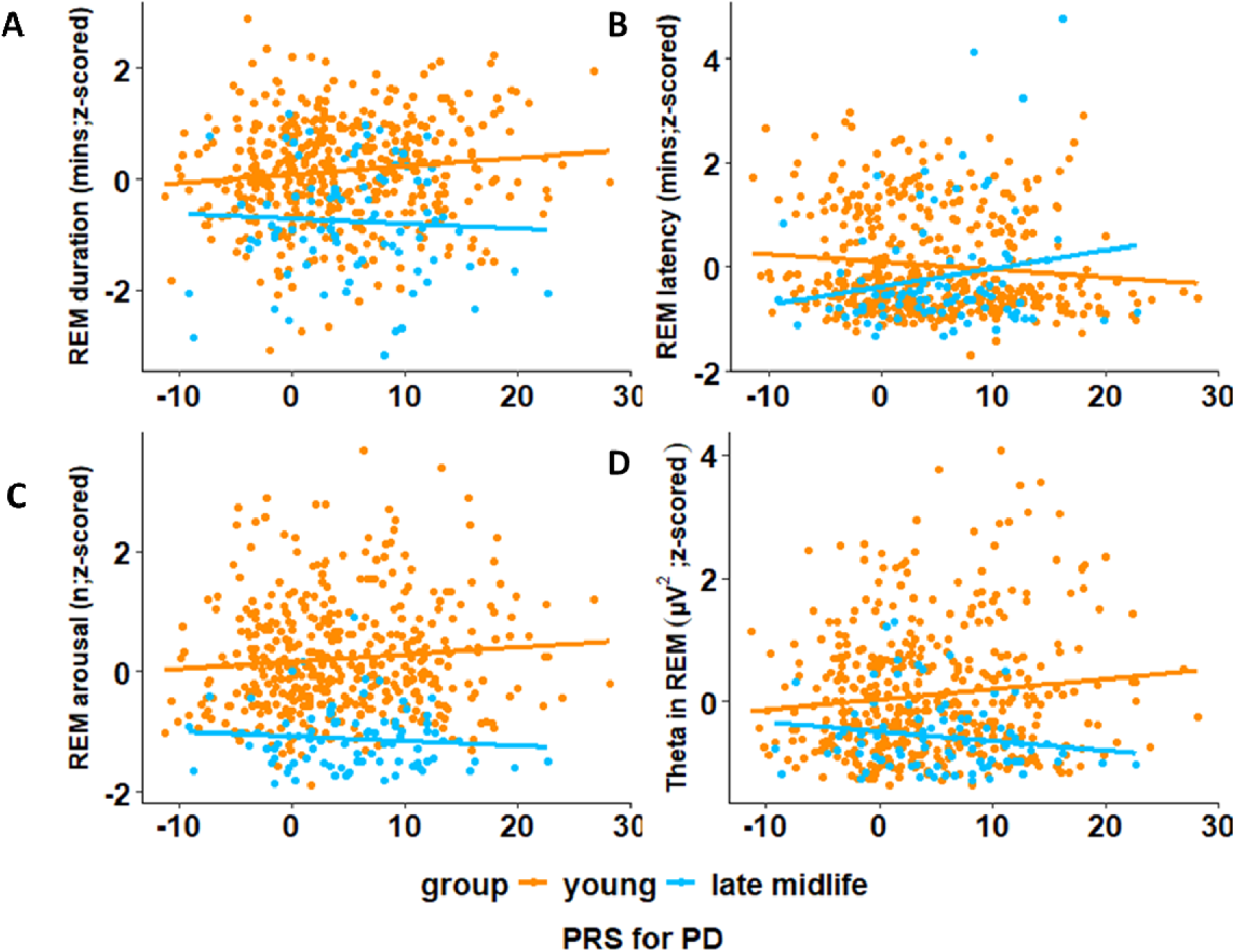
Age related aassociations between PRS for PD and baseline night sleep metrics for young and late-midlife sub-samples (A-D). The association between (A) REM duration and (D) theta in REM during baseline night and PD PRS were significant. Refer to main text **Table 3** for complete statistical outputs of GAMLSSs.

**Figure 4:**
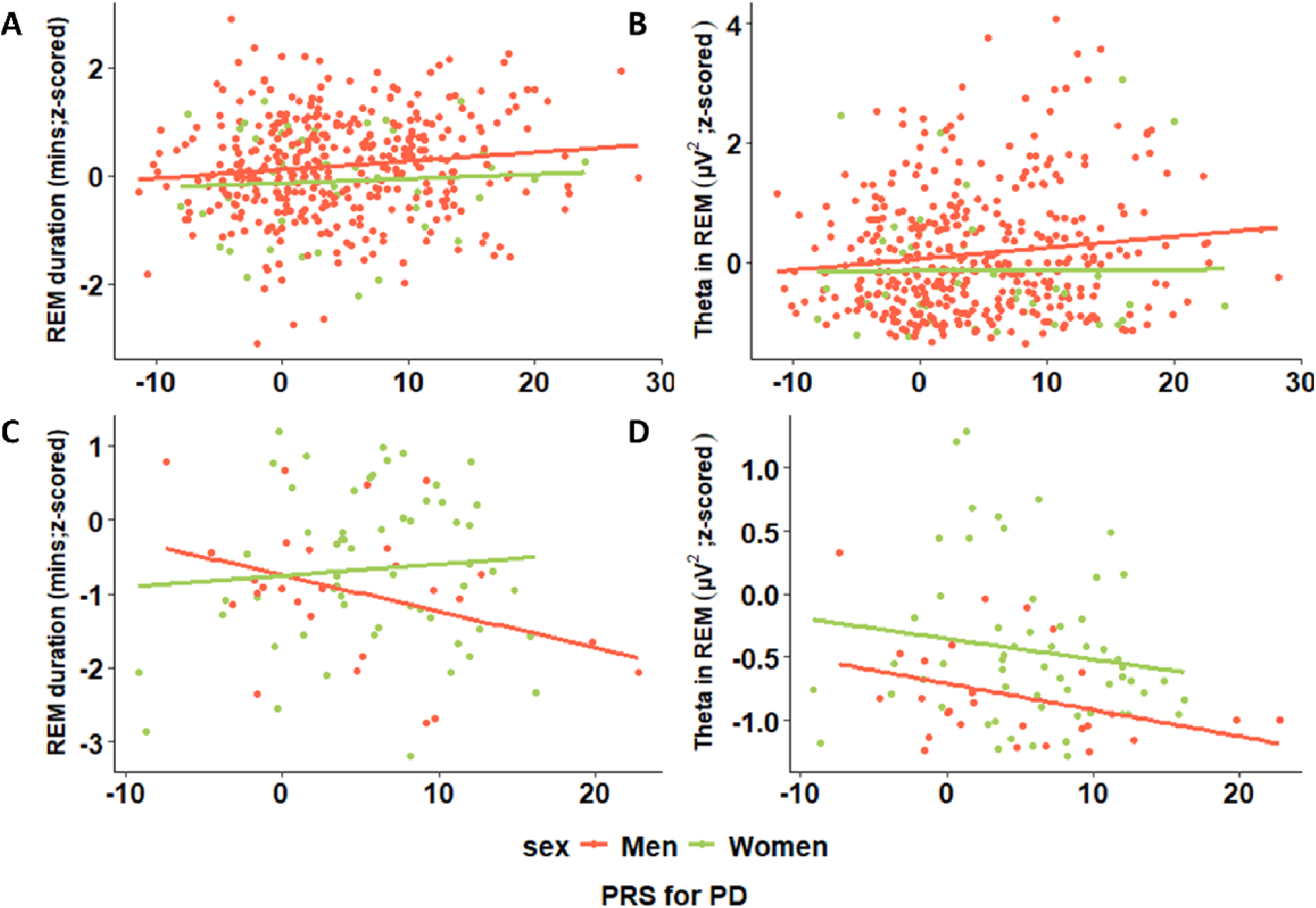
Sex specific aassociations between PRS for PD and baseline night sleep metrics for the young and late-midlife sub-samples. In the young sub-sample (upper panel), GAMLSS yielded no sex specific association for (A) REM duration and (B) theta in REM with PD PRS. In the late midlife subgroup (lower panel), GAMLSS yielded significant sex specific association for (C) REM duration with PD PRS but not for (D) theta in REM. Refer to supplementary **Table S1 and S2** for complete statistical outputs of GAMLSSs.

**Table 3.**
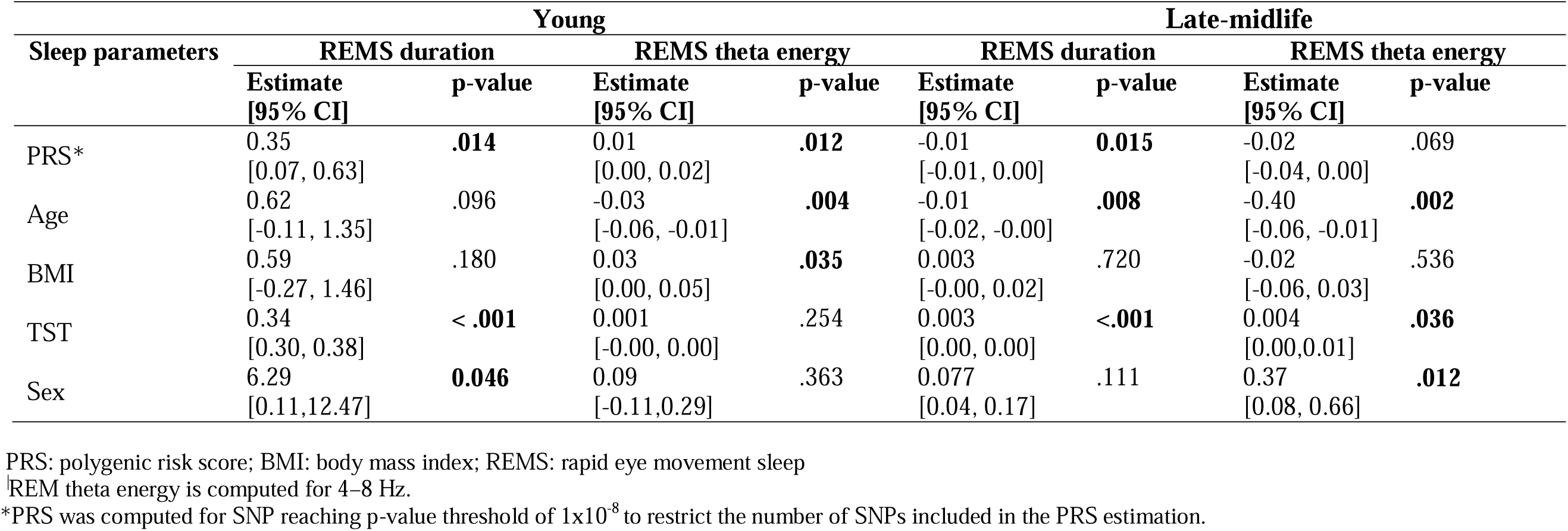
Results derived from regression analysis when testing for associations between sleep parameters and PRS values computed for Parkinson’s disease risk in healthy younger (N=345) and late-midlife sub-sample (N=85).

| Sleep parameters | Young |  |  |  | Late-midlife |  |  |  |
| --- | --- | --- | --- | --- | --- | --- | --- | --- |
|  | REMS duration |  | REMS theta energy |  | REMS duration |  | REMS theta energy |  |
|  | Estimate<br>[95% CI] | p-value | Estimate<br>[95% CI] | p-value | Estimate<br>[95% CI] | p-value | Estimate<br>[95% CI] | p-value |
| PRS* | 0.35<br>[0.07, 0.63] | **.014** | 0.01<br>[0.00, 0.02] | **.012** | -0.01<br>[-0.01, 0.00] | **0.015** | -0.02<br>[-0.04, 0.00] | .069 |
| Age | 0.62<br>[-0.11, 1.35] | .096 | -0.03<br>[-0.06, -0.01] | **.004** | -0.01<br>[-0.02, -0.00] | **.008** | -0.40<br>[-0.06, -0.01] | **.002** |
| BMI | 0.59<br>[-0.27, 1.46] | .180 | 0.03<br>[0.00, 0.05] | **.035** | 0.003<br>[-0.00, 0.02] | .720 | -0.02<br>[-0.06, 0.03] | .536 |
| TST | 0.34<br>[0.30, 0.38] | **< .001** | 0.001<br>[-0.00, 0.00] | .254 | 0.003<br>[0.00, 0.00] | **<.001** | 0.004<br>[0.00,0.01] | **.036** |
| Sex | 6.29<br>[0.11,12.47] | **0.046** | 0.09<br>[-0.11,0.29] | .363 | 0.077<br>[0.04, 0.17] | .111 | 0.37<br>[0.08, 0.66] | **.012** |

PRS: polygenic risk score; BMI: body mass index; REMS: rapid eye movement sleep
REM theta energy is computed for 4–8 Hz.
\*PRS was computed for SNP reaching p-value threshold of $1 \times 10^{-8}$ to restrict the number of SNPs included in the PRS estimation.

The GAMLSS per age sub-group further showed that the significant association between REM theta energy and PRS for PD was positive in the younger sub-sample (p=0.012), i.e. higher REM intensity is related to higher PD risk, while, though it was negative, it was not significant as a main effect in the late-midlife sub-sample and only reach statistical trend values (p=0.07; **Table 3**, **Fig. 3B**). Moreover, in contrast to REM duration, including sex in the GAMLSS of the young and late midlife sub-sample did not reveal sex or sex-by-PRS interactions (**Suppl. Table S1**, **Fig. 4B&D**). Therefore, as with REM duration, REM intensity as measured by REM theta energy is associated with PD PRS in an age dependent manner with the direction of association changing with age.

### Specificity - Association with other sleep metrics

To establish the robustness of our findings we recomputed the PRS for PD using a larger number of common variants. The GAMLSS using REM duration and theta energy in REM yielded the same statistical outputs including slightly more variants, but not many more nor all genetic variants (**Suppl. Fig. S2**) supporting the need for a narrow/specific PRS for PD for association to emerge. In addition, there was no association with a polygenic prediction of an individual physical trait with which REM sleep is not expected to be associated (height) showing that our results cannot be obtained with any polygenic computations (**Suppl. Fig. S3**).

To further ascertain specificity of our findings, we considered additional sleep metrics: 1) REM percentage; 2) REM delta energy; 3) REM alpha energy; 4) REM sigma energy; 5) and Slow wave energy (SWE) during NREM sleep (see methods) (**Suppl. Fig. S4).** GAMLSS analyses over the entire sample for each of these metrics yielded significant age-group by PRS for PD interaction for REM sleep percentage (p<0.001) while controlling for body mass index (BMI) and REM duration (see methods) (**Suppl. Table S2**). These findings substantiate the previous results with REM duration. GAMLSS with REM delta energy, REM alpha energy, REM sigma energy or SWE in NREM sleep showed no statistically significant association. GAMLSS per sub-group yielded similar findings for REM percentage as for the primary analysis on REM duration. (**Suppl. Fig. S5**).

## Discussion

Although it is well accepted that sleep is altered over PD neuropathology – with RBD almost inescapably predicting the future development of Parkinsonism^6^ - how the association between REM sleep and PD emerges over the lifetime is not established. Here, we find that the polygenic liability, or PRS, for PD is related to REM sleep metrics - mainly REM duration and theta energy in REM. The associations were isolated in healthy young adults aged 31 or less, i.e. 2 to 5 decades before typical age of diagnosis, as well as in healthy late-middle aged individuals age 50 to 69y. Higher PRS for PD was associated with longer REM sleep duration and more intense REM sleep, as assessed by the overnight REM theta energy, in the younger subsample. By contrast, higher PRS for PD was correlated with shorter REM sleep duration, potentially specifically in men, and was qualitatively – though not statistically – associated with lower REM theta energy. The negative control analysis, using polygenic prediction of height, together with the fact that no association was found between PRS for PD and SWE or other frequency bands of REM sleep (except for delta energy), supports that the associations are stronger and/or specific to REM sleep duration and theta energy. The effect size we uncover are small, as in any such studies, yet their cumulative effect over the years could have important implications underlying a true contribution of sleep to PD neurobiology. In addition, the associations are found based on electrophysiology measures of sleep, which contrasts to prior studies that could only access coarser non-electrophysiological phenotyping consisting of e.g. sleep questionnaires or actigraphy alone^25^.These findings add to the current literature and may isolate the earliest and potentially core links between REM sleep and PD-related biology.

Sleep disturbances often precede by decades the onset of motor symptoms in PD^26^. Our findings suggest that these alterations may first take the form of a longer overnight REM sleep and of an increase in REM theta energy, which gathers the overnight power of the most typical oscillatory mode of REM sleep. The somewhat stronger initial manifestation of REM sleep could suggest that the biology of prodromal PD leads to larger capacity for REM sleep-related processes such as memory consolidation, emotional processing, brain development, and/or dreaming^27^ in those young adults with higher PD genetic liability. The biology of individuals with higher PRS for PD may lead to a stronger expression of REM, inherently - for an equivalent brain function during wakefulness - or in response to wakeful events previously experienced by the young individuals. Both options could be tested by linking electrophysiology metrics acquired during wakefulness to PRS for PD in young adult individuals. One could further test whether the PRS-for-PD-dependent manifestation of REM sleep is associated with difference in sleep-dependent memory consolidation or emotional processing.

Our results further support that later over the lifetime, while still healthy, the same associations switch to a shorter REM sleep duration and a lower REM theta energy in those with a higher genetic liability for PD. This is in line with observations that PD patients present reduced REM sleep duration^28^. In a longitudinal study involving 2770 healthy late-midlife men, lower REM percentage were found to be associated with an increased risk of developing PD^29^. It does not match however the report that patients with PD-RBD may exhibit higher theta spectral energy during REM sleep^30^ and that PD patients typically show a sustained increase in high-theta/alpha frequencies (7.8-10.5 Hz) early in the sleep period ^31^.

The reasons for the opposite age-related associations cannot be identified through our cross-sectional study. A first potential explanation may, however, be related to the high oxidative metabolism and catecholamine activity imposed by REM sleep^32^. Small structures of the brainstem such as the SN and LC are known to be particularly sensitive to oxidative insult and high cytosolic catecholamine concentration. This would be in fact the reason why both structures show progressive high level of neuromelanin over the lifespan as it is considered that neuromelanin serves to shield the cells from toxic effects of redox active metals, toxins, and excess of cytosolic catecholamines^33^. The LC is a nucleus crucial for the transitions between NREM and REM sleep^34^ while dysfunction and degeneration of noradrenergic neurons in the LC are linked to RBD^35^ and reduced arousal. The SN also contributes to REM regulation^36^ and is central to PD motor symptoms^37^. We speculate therefore that those young individuals with higher PRS which show high REM duration and intensity based on our data, would show a quicker alteration of structures such as the SN and LC leading to progressive reduction in REM duration and intensity.

A second plausible explanation, that is not mutually exclusive with the first one, is that the opposite age-related association in REM sleep – PRS association it may be due to the progressive accumulation of α-synuclein and Lewy bodies, which is presumably quicker in individuals with higher PRS for PD. The earliest brain α-synuclein deposits are observed long before the onset of motor symptoms in brainstem structures of the extended medulla and pons (Braak stage 1 and 2, ^38^), among which the LC. Likewise, VTA dopaminergic neurons lesion in rats reduced total NREM and REM sleep duration during the sleep phase^39^. We posit therefore that α-synuclein aggregates in the LC and maybe in the VTA, and in the SN, contributes to the association we find between REM duration and PRS for PD.

The hippocampus is the source of ripple oscillations which are essential to the interplay between the limbic system, the thalamus and the cortex that contributes to memory consolidation^40^. Ripples cannot be directly detected *per se* on the EEG which arises mostly from the activity of cortical neurons, but they contribute to theta cortical oscillations in the cortex^41^. In addition, lesions of the VTA dopaminergic neurons in rats were found to lead to suppression of EEG theta rhythm frequency during both wakefulness and REM sleep, suggesting that midbrain dopaminergic neurons contribute to hippocampal theta activity^42^. The changes in theta oscillation energy could therefore arise from the putative α-synuclein aggregates in the hippocampus or in the VTA.

In addition, later, when α-synuclein or neuromelanin is eventually shed in the extracellular space upon neuronal death, the neuroinflammation response and/or modulation of glymphatic system by sleep state could further influence PD process^10, 33^. It is therefore also possible that early REM sleep alteration contribute to the neurobiology of PD and to the progressive increase in oxidative insults and misfolded α-synuclein burden^6, 10^, which would in turns alter sleep regulation and oscillations.

The cross-sectional nature of our study precludes causal inferences and is not its only limitation. The exclusion criteria for participant selection were rigorous and not common for large genetic studies. As poor sleep, daytime sleepiness and REM without atonia (RSWA) can occur in the prodromal phase of PD, by excluding such participants we excluded some of the highest-risk individuals, especially among older participants. The goal of the study was to examine REM sleep and PD biology in the absence of comorbidities that could bias associations, our approach reduces confounding and strengthens internal validity but limits generalizability and may attenuate age-related differences, highlighting the need for future studies including individuals with diverse health conditions. As mentioned earlier, sex was included as covariate in all analyses, but our sample predominantly consisted of men in younger sub-sample, and of mainly women in older sub-sample, so potential sex differences could not be properly studied here. The sex difference in the relationship between PRS for PD and REM duration that we report in the late midlife group has therefore to be taken with caution. Yet a qualitative consideration of the data collected in women, do not indicate sex difference in our analyses of interest in the younger group. This warrants future investigation including samples balances for sexes, as women present different sleep characteristics and difference in the age-related changes in sleep^43, 44^ in addition to the fact that they are less prone to PD^45^.

Although 15% of PD patients have a positive family history, and 5-10% of cases follow a Mendelian inheritance pattern, the aetiology of PD is multifactorial. It is strongly influenced by environmental factors, with age being the most important risk factor^3^. We therefore emphasize that a polygenic risk score (PRS) for Parkinson’s disease (PD) is not intended for individual prediction. Instead, PRS are used to link the underlying biology of the disease—since polygenic risk is inherently biology-grounded—to relevant phenotypes, including that observed in young adults (or even earlier). Accordingly, we present compelling evidence of an early association between REM sleep and PD-related biology, warranting future investigations to provide the functional understanding of the brain mechanism at stake.

Despite ongoing development of potential neuroprotective agents such as GLP-1 agonists, calcium channel blockers, and urate, their efficacy in modifying disease progression remains unproven^5^. We argue that the detection of early association between REM sleep and PD through PRS could improve the early prediction of PD risk and provide novel intervention targets. However, as PRS is a risk indicator, a longitudinal study showing that REM sleep changes in at risk individuals in association with onset of PD symptoms later in life would be a necessary validation step. The sleep-based interventions could take the form of cognitive behavioral therapy, light therapy, LC stimulation^46^, or pharmacological approaches targeting dopamine or norepinephrine modulation^47^ and would need to be tested in large scale interventional studies. Studies shows for instance that rotigotine, a transdermal dopamine agonist increases REM sleep stability^48^. Whether it bears positive outcomes for PD is not known.

In conclusion, our analysis involving the detailed electrophysiological phenotyping of a relatively large sample size of 518 healthy individuals showed that PRS for PD is related to REM sleep metrics, specifically its duration and theta energy. The relationship between PRS and sleep switches from positive to negative across younger and late-midlife groups. Our findings suggest that the association between PRS and quantitative REM sleep metrics in late-midlife group is moderated by sex, which may be related to the sex-effects in PD prevalence^2^. Quantitative sleep measures could help in the early detection of the underlying neurodegenerative process leading to several neurological diseases including PD, 2 to 5 decades prior to typical symptom onset. The early identification of at-risk individuals for developing PD would allow evaluation of possible sleep targeted therapies as well as several neuroprotective agents.

## Methods

### Participants

The study sample comprised of 538 (age: 28.09±14.1y, 104 females) healthy Caucasian participants recruited from the local French-speaking community as part of 7 different multi-modal cross-sectional studies. One study contributed to most of the younger sample and included 359 young men to maximize genetic uniformity in what was originally a genetic study^19^. Remaining data of the younger sample (N=92; 45 women) was collected over 5 different studies^49–53^. In contrast, data from late middle-aged individuals (N=87; 59 women) were collected as part of a single study^18^. All studies collected quantitative sleep parameters and blood samples to assess PRS for PD (Figure 1). Participants with poor subjective sleep quality, daytime sleepiness, sleep disorders (including significant sleep apnoea, parasomnia, RBD and RSWA), addiction, cognitive impairment, or taking any medications impacting the CNS were excluded. Further details are provided in **Suppl. file 1.**

The study procedures were approved by the Ethics Committee of the Faculty of Medicine (University of Liège, Belgium). All participants gave their written signed informed consent prior to their participation in the study and received financial compensation. The study was conducted in accordance with the World Medical Association International Code of Medical Ethics (Declaration of Helsinki) for experiments involving humans.

Out of 538 (451 young and 87 old), 14 participants were excluded due to incomplete baseline data, 14 due to lack of genetic data and 6 outliers with ± 5S.D resulting in a final sample of 518 (433 young and 85 old) participants. The characteristics of the final participant sub-sample are reported in **Table 1**.

### Sleep protocol, EEG acquisitions and processing

The in-lab EEG recordings of sleep collected across 7 studies^17,18,49–53^ included baseline EEG of night-time sleep at habitual sleep times following at least one week of regular sleep-wake schedules monitored by actigraphy. More details (in brief) are provided in **Suppl. file 1.**

Scoring of sleep stages was performed using a validated automatic algorithm (ASEEGA, PHYSIP, Paris, France) in 30-s epochs^54^, according to 2017 American Academy of Sleep Medicine criteria, version 2.4. An automatic artefact and arousal detection algorithm with adaptative thresholds^55^ was further applied and artefact and arousal periods were excluded from subsequent analyses. Power spectrum was computed for each channel using a Fourier transform on successive 4-s bins, overlapping by 2-s, resulting in a 0.25 Hz frequency resolution. The night was divided into 30 min windows, from sleep onset, defined as the first NREM2 (N2) stage epoch, until lights-on. Averaged power was computed per 30 min bins, adjusting for the proportion of rejected data, and subsequently aggregated in a sum separately for REM and NREM sleep^56^. Thus we computed slow wave energy (SWE) - cumulated power in the delta frequency band during N2 and N3 sleep stages, an accepted measure of sleep need^56^, and similar to that we computed the cumulated theta (4-8Hz) power in REM sleep. We then computed the cumulated power over the remaining EEG bands, separately for NREM and REM sleep: alpha (8-12Hz), sigma (12-16Hz), beta (16-25Hz), theta (4-8Hz) and delta (0.5-4 Hz) bands. As the frontal regions are most sensitive to sleep–wake history^56^, we considered only the frontal electrodes (mean over F3, Fz, and F4), as well as to facilitate interpretation of future large-scale studies using headband EEG, often restricted to frontal electrodes.

Our analyses focused on four sleep metrics to limit issues of multiple comparisons while spanning the most important aspects of REM sleep EEG: 1) REM duration; 2) REM latency; 3) number of arousals during REM sleep; 4) cumulated theta power during REM sleep. To ascertain specificity of findings we also considered 1) REM percentage, which reflect the overall architecture of sleep rather than only the duration of REM; other frequency band of the EEG during REM, i.e. 2) REM alpha energy, 3) REM sigma energy, as well as 4) REM delta energy, as the definition of REM frequencies during REM varies across publication; and finally 5) SWE during NREM sleep, the dominant oscillatory mode of NREM sleep, considered to be tightly related to the need for sleep^56^.

### Genotyping, quality control and imputation

The blood samples or buccal swabs were collected and stored at -20°C within few hours until DNA extraction. The genotyping was performed at different time points using the Illumina Infinium OmniExpress-24 BeadChip arrays (Illumina, San Diego, CA) based on Human Build 37 (GRCh37) at Genomics platform of Liège GIGA institute. All the study participants were of European ancestry. Established quality control (QC) procedure was performed using PLINK^57^ (http://zzz.bwh.harvard.edu/plink/). In brief, the SNPs were excluded as follows: >10% missing genotypes, <95% call rate, minor allele frequency (MAF) below 0.01, out of Hardy-Weinberg equilibrium (p-value <10^-4^ for the Hardy-Weinberg test). SNPs on 23rd chromosome as well as ambiguous SNPs (A-T, T-A, C-G, G-C) were excluded as well. The data was matched for deviation with European ancestry using 1000 Genomes Project dataset (1KGP, https://www.internationalgenome.org). Imputation was conducted using the Sanger imputation server (https://imputation.sanger.ac.uk/) based on the Haplotype Reference Consortium (r1.1) as reference panel and using EAGLE2 pre-phasing algorithm. The detailed data processing and analysis for young and late-midlife sub-sample is as described previously in^17, 19^. We finally ended with 7,165,614 SNPs.

### Polygenic risk score calculation

Polygenic risk score (PRS) analyses can be used to assess the genetic liability of an individual for a phenotype by calculating the weighted sum of risk alleles effect size identified in genome-wide association studies. In the current study, a PRS for Parkinson’s disease based on summary statistics from the recent meta-analysis GWAS for Parkinson’s disease of European ancestry was calculated for each participant^21^.The standardization and quality control of GWAS summary statistics was performed by MungeSumstats, a Bioconductor R package^58^. In the process, the summary statistics was pruned to align reference alleles to build GRCh37, remove multiallelic variants, and adjust weights for the appropriate reference alleles. The PRS was then generated using SBayesR algorithm implemented in GCTB software. The approach assumes that the SNP effects are drawn from mixtures of distributions with the key metrics defining these genetic architectures estimated through Bayesian frameworks. To derive PRSs from GWAS effect estimates of SNPs, SBayesR essentially uses Bayesian linear mixed model and the reference linkage disequilibrium (LD) correlation matrix. In our analysis, we used banded LD matrix, derived from ∼1.1 million HapMap3 SNPs in 50,000 unrelated UK Biobank participants of European ancestry, to improve prediction accuracy as recommended by the authors of GCTB. Applying SBayesR with default parameters to GWAS summary statistics, we obtained updated effect sizes for 1,104,064 SNPs, which were then used to calculate the PRS. We then used p-value thresholding through PLINK to include only the SNPs reaching stringent GWAS significance (p-value <10^-8^) to restrict the number of genetic markers. This stringent threshold was pre-specified as our main analytic approach. Analyses at alternative thresholds were conducted to examine the stability of associations under less conservative variant inclusion criteria.

### Height as a negative control

Based on the current available literature on sleep biology, we assumed absence of any a priori association between the sleep phenotypes and a genetic liability for height. Therefore, we conducted an analysis of polygenic scores estimated for height as a negative control, performing the same GAMLSS analyses as we did for liability to PD.

### Statistical analysis

All analysis was conducted within the R environment (version 4.1.3) (R Development Core Team, 2017). We employed generalized additive models for location scale and shape (GAMLSS)^24^ to individually assess the associations between four sleep metrics of interests (REM duration, Theta in REM, REM latency and No. of arousals in REM), as dependent variable, and the PRS values for PD as an independent variable. GAMLSS offers a wide variety of family of distributions for model fitting^59^ and are considered better than GLM or GAM approaches (**Suppl. Fig. S6)**. In our analysis only the location parameter (μ) is modeled as a function of the covariates. The estimates (β) and confidence interval (CI) are reported in log scale.

Individual values in the dataset were considered outliers if >5SD from the mean and excluded from analyses. For fitting GAMLSS models, family was selected based on data distribution using fitDist function. Further, models were selected based on the goodness-of-fit values among GAMLSS models and based on the Q-Q plot. Sex, BMI, total sleep time (TST) or REM duration were included as covariates. Prior to the GAMLSS analysis, influential outliers were also screened using worm plot, a detrended Q-Q plot, helpful for checking model fit and comparing the fit of different models. We have provided a representational model fitting diagnostic plot as a supplementary figure (**Suppl. Fig. S6)**. Benjamini & Hochberg False Discovery Rate (FDR) correction was applied to the four pre-specified primary sleep metrics. Analyses of additional sleep metrics and alternative PRS thresholds were performed as exploratory robustness checks. These supplementary results are therefore interpreted with caution and without further multiple testing correction, as their role is to support the consistency of the primary findings. Sleep metrics were standardized using a Z-transformation for plots.

We computed a priori sensitivity analysis to get an indication of the minimum detectable effect size in our main analyses given our sample size. Based on G*Power 3 (version 3.1.9.4)^60^ (https://puneet-talwar.shinyapps.io/PosthocPowerShiny/) taking into account a power of .8, an error rate α of .0125 (corrected for 4 tests), a sample size of 518 allowed us to detect small effect sizes f2 >.036 (confidence interval: .018 –.09; R² > .035, R² confidence interval: .018 –.082) within a linear multiple regression framework including 2 tested predictor (PRS, age) and 4 other covariates (sex, BMI, total sleep time (TST) or REM duration). Prior sensitivity analysis for younger and late midlife sub-groups is provided in the supplementary file.

## Acknowledgements

PT is supported by the EU Joint Programme Neurodegenerative Disease Research (JPND) IRONSLEEP project (FNRS reference: PINT-MULTI R.8011.21). FC, CP and CBastin and GV are supported by the Fonds de la Recherche Scientifique - FNRS-Belgium. The study was supported by the Wallonia-Brussels Federation (Actions de Recherche Concertées - ARC—09/14-03,17/27-09), WELBIO/Walloon Excellence in Life Sciences and Biotechnology Grant (WELBIOCR-2010-06E), FNRS-Belgium (FRS-FNRS, F.4513.17 and T.0242.19, T.0238.23, J. 0222.20 and 3.4516.11), Fondation Recherche Alzheimer (SAO-FRA 2019/0025,2022/0014), University of Liege (ULiege), Fondation Simone et Pierre Clerdent, European Regional Development Fund (Radiomed project), Fonds Leon Fredericq, EU JPND program (IRONSLEEP project - PINT-MULTI R.8011.21). The authors also thank Sarah Chellappa, Annick Claes, Catherine Hagelstein, Gregory Hammad, Brigitte Herbillon, Patrick Hawotte, Mathieu Jaspar, Sophie Laloux, Erik Lambot, Benjamin Lauricella, André Luxen, Julie Ly, Christelle Meyer, and Eric Salmon for their help over the different steps of the study. This work was conducted at the GIGA-In Vivo Imaging platform of ULiège, Belgium.

## Author contributions

PT, PM and GV designed the experiment. PT, NM, EK, VM and CB analysed the data. PT and GV wrote the paper. MZ, FC, CD, CB, CBastin and CP provided support for data acquisitions, data analyses, administrative and/or financial aspects of the study. All authors edited the draft manuscript and approved its final version.

## Competing Interests Statement

Christian Berthomier is an owner of Physip, the company that analysed the EEG data. This ownership and the collaboration had no impact on the design, data acquisition, results and interpretations of the findings. The other authors declare that no competing interests exist.

## Data availability

The data and analysis scripts supporting the results included in this manuscript are publicly available via https://gitlab.uliege.be/CyclotronResearchCentre/Public/xxx (to be done following peer reviewing and upon acceptance for publication/and editor request). The following shiny app developed by PT (https://puneet-talwar.shinyapps.io/GAMLSSToolbox/) was also used for the GAMLSS analysis. We used Matlab scripts for EEG and MRI data processing, while we used R studio for statistical analyses. Researchers willing to access the raw data should send a request to the corresponding author (GV). Data sharing will require evaluation of the request by the local Research Ethics Board and the signature of a data transfer agreement (DTA).

## Notes

### Competing Interest Statement

Christian Berthomier is an owner of Physip, the company that analyzed the EEG data. This ownership and the collaboration had no impact on the design, data acquisition, results and interpretations of the findings. The other authors declare that no competing interests exist.

### Summary of Updates

The revised paper includes more data (~100 young participants) in the analyses, with about 50 women. Substantial revision were made to the methods and discussion sections.

