## Supplementary file for "Age-related differences in the association between REM sleep and the polygenic risk for Parkinson’s disease"

**Online Supplementary Information**

Puneet Talwar^1*^, N. Mortazavi^1^, Ekaterina Koshmanova^1^, Vincenzo Muto^1^, Aurora Gasparello^1^, Christian Degueldre^1^, Christian Berthomier^2^, Fabienne Collette^1^, Christine Bastin^1^, Christophe Phillips^1^, Pierre Maquet^1,3^, Zubkov Mikhail^1^, Gilles Vandewalle^1*^

1. GIGA-Institute, CRC-Human Imaging, University of Liège, Belgium
2. Physip, Paris, France
3. Department of Neurology, CHU of Liège, Liège, Belgium

***Address for Correspondence:**

Puneet Talwar & Gilles Vandewalle, PhD

GIGA-Institute, CRC-Human Imaging

Bâtiment B30, Université de Liège, 4000 Liège, Belgium

Phone : +32-4366-2316

### Supplementary Methods

### *Participants*

In brief, for all study groups participants with body mass index (BMI) greater than 27 kg/m²; a history of psychiatric conditions or severe brain injury; addiction; chronic use of medication affecting the central nervous system (CNS); smoking, excessive alcohol consumption (more than 14 units per week), or high caffeine intake (more than 3 cups per day); shift work in the past year; trans-meridian travel in the past 3 months; moderate to severe subjective anxiety and depression, as assessed by the Beck Anxiety Inventory^1^ (BAI; score greater than 16) and Beck Depression Inventory II^2^ (BDI; score greater than 19), respectively were excluded from the study. Participants with poor sleep quality, as measured by the Pittsburgh Sleep Quality Index^3^ (PSQI; score greater than 7), excessive daytime sleepiness, as indexed by the Epworth Sleepiness Scale^4^ (ESS; score greater than 15), or significant sleep apnea (apnea-hypopnea index greater than 15 per hour, according to the 2017 American Academy of Sleep Medicine criteria, version 2.4) were also excluded based on an in-lab screening night of polysomnography. For late-midlife participants, the exclusion criteria were relaxed for BMI (> 30 kg/m2) and excessive consumption of caffeine (>4 cups/day). While most participants scored normal ESS values (≤11), 37 participants had scores ranging from 12 to 15, corresponding to moderate daytime sleepiness.

### *Sleep protocol*

### Most of the data of the young participants (N=344) were collected as part of the same study (REF) during individual sleep-wake cycle was strictly monitored. In brief, for three weeks prior to the in-lab experiment, participants adhered to a regular sleep schedule based on their usual sleep times (within ±30 minutes for the first two weeks and within ±15 minutes for the final week; no daytime napping), verified through actigraphy data (Actiwatch 4, CamNtech, Cambridge, UK). They were also instructed to refrain from unusually intense physical exercise for the last 3 days of the fixed-schedule circadian entrainment. Immediately before the experiment, participants underwent a urine drug test (Multipanel Drug Test, SureScreen Diagnostics Ltd) and completed an adaptation night following their habitual sleep/wake schedule. During this night, a full polysomnography was recorded to check for sleep-related breathing disorders or periodic limb movements. On the second day, participants left the lab in the morning with instructions not to nap, which was verified using actigraphy data. They returned to the lab at the end of the second day (3.5 hours before their scheduled lights-off) and had a baseline night of sleep (BAS) in complete darkness, centered on the average sleep midpoint of the preceding week. This study focuses solely on the baseline night of sleep.

### All the data of the late midlife participants (N=85) were collected as part of the same study^5^ which included a slightly different protocol. As part of the screening process, they first completed an in-lab adaptation and screening night to familiarize with the new environment. Then, for 7 days before the baseline night, participants adhered to a regular sleep-wake schedule (within ±30 minutes; no daytime napping) according to their habitual bed and wake-up times, verified using sleep diaries and wrist actigraphy (Actiwatch©, Cambridge Neurotechnology, UK). They were also instructed to refrain from unusually intense physical exercise for the last 3 days of the fixed-schedule circadian entrainment. Their habitual sleep was then recorded in complete darkness using EEG. The additional dataset of the young individuals (N=89) was collected following the same procedure as for the late midlife participants in 5 different studies (a week between adaptation and data collection nights)^6^.

***EEG acquisitions***

Polysomnographic sleep data of young participants were acquired using either V-Amp 16 amplifiers (Brain Products, Germany)^7-10^, a QuickAmp-72 (Brain Products, Germany)^11^ or a N7000 amplifiers (EMBLA, Natus Medical Incorporated, Planegg, Germany)^12^. The electrode montage consisted of at least 9 EEG channels (F3, Fz, F4, C3, Cz, C4, Pz, O1, O2; reference to right mastoid), 2 bipolar EOGs, 2 bipolar EMGs, and 2 bipolar ECGs. For late-midlife group, participants sleep EEG data were acquired using N7000 amplifiers (EMBLA, Natus Medical Incorporated, Planegg, Germany) with 10 EEG channels (F3, Fz, F4, C3, Cz, C4, P3, Pz, P4, O1, O2), 2 bipolar EOGs, 2 bipolar EMGs, and 2 bipolar ECGs. For both age groups, acquisition on the screening night of sleep also included respiration belts, oximeter and nasal flow, 2 electrodes on one leg, but included only Fz, C3, Cz, Pz, Oz, and A1 channels. Baseline EEG data were re-referenced off-line to average mastoids.

ASEEGA, a validated automatic algorithm, was used for the scoring of sleep stages. ASEEGA is validated for sleep scoring in younger and older individuals as well as in several sleep disorders^13^^,^ ^14^. This includes REM sleep detection where specificity and sensitivity were for instance reported to be respectively 95% and 89%. Our group has previously shown that the consensus for REM detection in older individuals was high between ASEEGA and 2 different sleep experts^15^.

### *Statistical analysis*

We computed a priori sensitivity analysis for the younger group given our sample size. Taking into account a power of .8, an error rate α of .05, a sample size of 433 allowed us to detect small effect sizes f^2^ =.032 (confidence interval: .02 –.09; R² > .031, R² confidence interval: .015 –.084) within a linear multiple regression framework including 2 tested predictor (PRS, age) and 4 other covariates (sex, BMI, total sleep time (TST) or REM duration). Additionally, we also computed a prior sensitivity for late midlife group. Taking into account a power of .8, an error rate α of .05, a sample size of 85 allowed us to detect medium-large effect sizes f^2^ =.187 (confidence interval: .09 –.56; R² > .157, R² confidence interval: .082 –.358) within a linear multiple regression framework including 2 tested predictor (PRS, age) and 4 other covariates (sex, BMI, total sleep time (TST) or REM duration). There is limited published data on the precise effect sizes for quantitative sleep metrics and genetic associations in late midlife adults. However, recent studies have used similar sample size in the analysis^16^ providing further support to the validity of our study results.

**Suppl.** **Table S1.** Results derived from regression analysis when testing for associations between sleep parameters and PRS values computed for Parkinson’s disease risk in young (N=433) and late-midlife cohort (N=85).

| **Sleep**  **parameters** | **Young** | | | | **Late midlife** | | | |
| --- | --- | --- | --- | --- | --- | --- | --- | --- |
|  | **REMS duration** | | **REMS theta Power** | | **REMS duration** | | **REMS theta Power** | |
|  | **Estimate**  **[95% CI]** | **p-value** | **Estimate**  **[95% CI]** | **p-value** | **Estimate**  **[95% CI]** | **p-value** | **Estimate**  **[95% CI]** | **p-value** |
| PRS*sex | 0.098  [-0.31, 0.40] | .81 | 0.01  [-0.01, 0.02] | .51 | -0.018  [-0.03, -0.00] | **.03** | -0.015  [-0.06, 0.03] | .476 |
| PRS^$^ | 0.26  [-0.22, 0.44] | .49 | 0.004  [-0.01, 0.01] | .78 | 0.002  [-0.01, 0.01] | .680 | -0.014  [-0.04, 0.01] | .322 |
| Age | 0.63  [-0.04, 0.59] | .09 | -0.034  [-0.03, -0.01] | **.003** | -0.010  [-0.02, -0.00] | **.033** | -0.038  [-0.06, -0.01] | **.003** |
| BMI | 0.59  [-0.12, 0.64] | .18 | 0.029  [0.00, 0.02] | **.03** | -0.006  [-0.02, 0.01] | .505 | -0.017  [-0.06, 0.03] | .440 |
| TST | 0.34  [0.13, 0.17] | **<.001** | 0.001  [-0.00, 0.00] | .25 | 0.004  [0.00, 0.00] | **<.001** | 0.003  [0.00,0.01] | **.046** |
| Sex | 5.84  [-0.56, 5.63] | .11 | 0.05  [-0.08, 0.12] | .67 | 0.034  [-0.10, 0.17] | .618 | -0.319  [-0.67, 0.03] | .074 |

PRS: polygenic risk score; BMI: body mass index; REMS: rapid eye movement sleep.

^$^PRS was computed for SNP reaching p-value threshold of 1x10^-8^ to restrict the number of SNPs included in the PRS estimation.

**Suppl.** **Table S2.** Results derived from regression analysis when testing for associations between REMS duration and PRS values computed for Parkinson’s disease risk in late-midlife cohort (N=85) segregated by sex.

| **Sleep**  **parameters** | **Men** | | **Women** | |
| --- | --- | --- | --- | --- |
|  | **Estimate**  **[95% CI]** | **p-value** | **Estimate**  **[95% CI]** | **p-value** |
| PRS^$^ | -0.016  [-0.01, -0.002] | **.005** | -0.001  [-0.01, 0.01] | .812 |
| Age | -0.014  [-0.01, 0.00] | **.05** | -0.013  [-0.02, -0.00] | **.017** |
| BMI | -0.019  [-0.02, 0.01] | .22 | 0.004  [-0.01, 0.22] | .656 |
| TST | 0.003  [0.00, 0.00] | **.003** | 0.004  [0.00,0.00] | **<.001** |

PRS: polygenic risk score; BMI: body mass index; REMS: rapid eye movement sleep.

^$^PRS was computed for SNP reaching p-value threshold of 1x10^-8^ to restrict the number of SNPs included in the PRS estimation.

**Suppl.** **Table S3.** Results derived from regression analysis when testing for associations between additional sleep parameters and PRS values computed for Parkinson’s disease risk in healthy younger (N=433) and late-midlife cohort (N=85).

| **REM percentage** | | | | | | |
| --- | --- | --- | --- | --- | --- | --- |
| **Study groups** | **All** | | **Young** | | **Old** | |
| **Model**  **parameters** | **Estimate**  **[95% CI]** | **p-value** | **Estimate**  **[95% CI]** | **p-value** | **Estimate**  **[95% CI]** | **p-value** |
| PRS*age group | **.01**  **[0.01, 0.02]** | **.001** | ̶ | ̶ | ̶ | ̶ |
| Age group | **-.01**  **[-0.07, 0.06]** | **.781** | ̶ | ̶ | ̶ | ̶ |
| PRS^$^ | -.01  [-0.02, -0.00] | .009 | 0.003  [0.00,0.01] | **.003** | -0.01  [-0.02, -0.00] | **.010** |
| Age | ̶ | ̶ | 0.003  [-0.00, 0.01] | .317 | -0.01  [-0.02, -0.00] | **.031** |
| BMI | .002  [-0.00,0.01] | .562 | 0.004  [-0.00, 0.01] | .291 | 0.003  [-0.01, 0.02] | .695 |
| Sex | .01  [-0.04,0.06] | .772 | 0.043  [-0.01, 0.10] | .123 | -0.09  [-0.19,0.01] | .070 |

PRS: polygenic risk score; BMI: body mass index; REMS: rapid eye movement sleep

^$^PRS was computed for SNP reaching p-value threshold of 1x10^-8^ to restrict the number of SNPs included in the PRS estimation

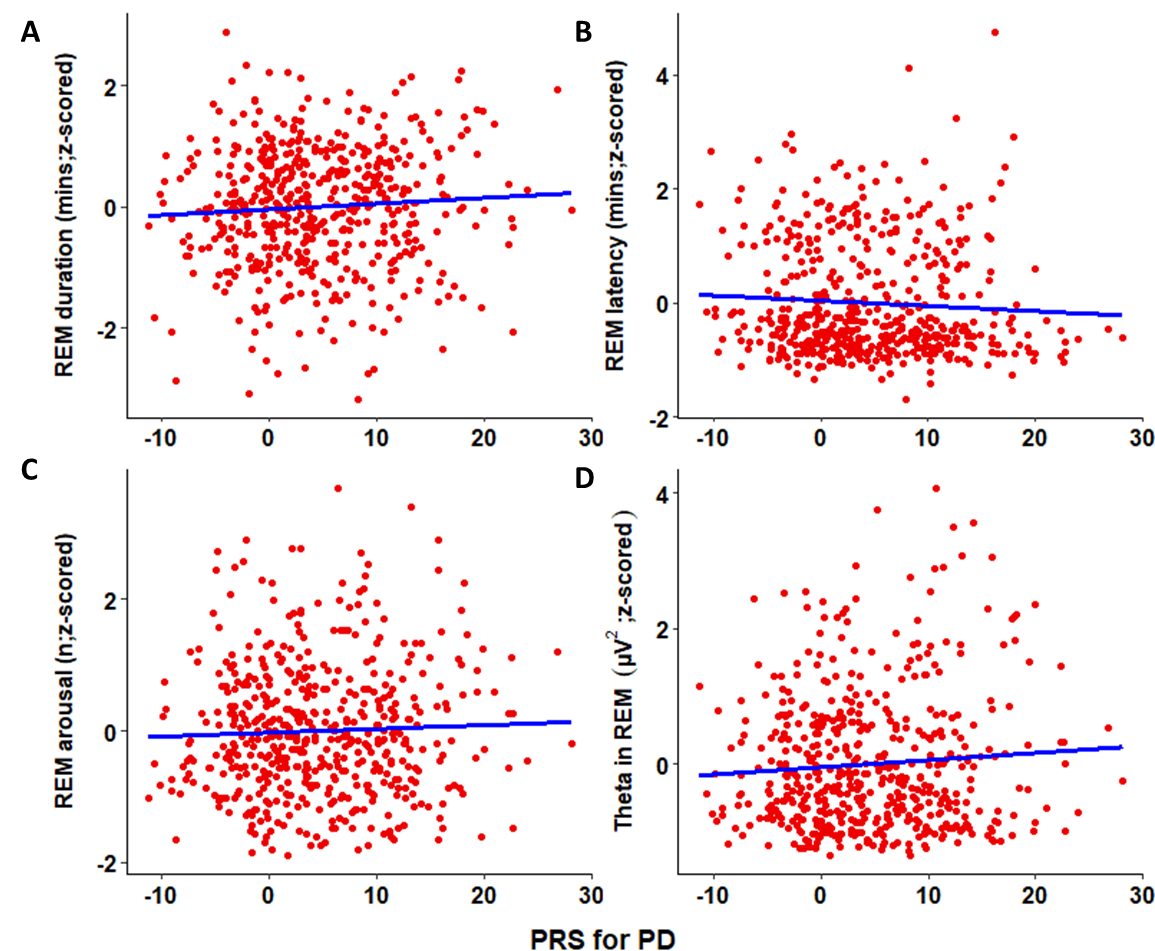

**Figure S1:** The associations between four sleep metrics during baseline night and PD PRS (*N* = 518). Spearman correlation *r* is reported for completeness; refer to main text **Table 2** for statistical outputs of GAMLSSs.

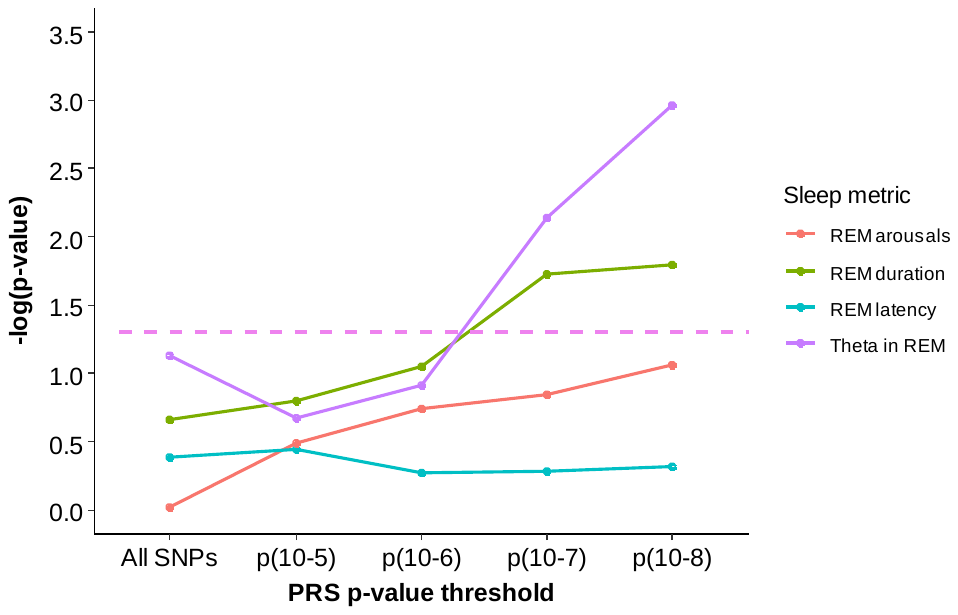

**Figure S2:** Associations between baseline night sleep metrics and PRS for PD computed using more SNPs. Statistical outcomes of GAMLSSs with four sleep metrics of interest versus PD PRS from conservative (p < 10^−8^) p value threshold to using all SNPs (N = 518). GAMLSSs are corrected for sex, BMI, and TST or REM duration. The negative log transformation of p values of the associations is presented on the vertical axis. Horizontal line (pink dashed line) indicates p values threshold of 0.05. These additional results show that progressively including less specific SNPs progressively weakens the associations (still significant using 10-^7^ threshold) which is coherent with a non-random association specific to PD biology. However, REM latency and arousal in REM did not reveal significant associations with PRS for PD at any p-value thresholds. Further, at lower p values threshold (10^-5^) and models including all SNPs in the PRS showed absence of significant association for all the sleep metrics of interest.

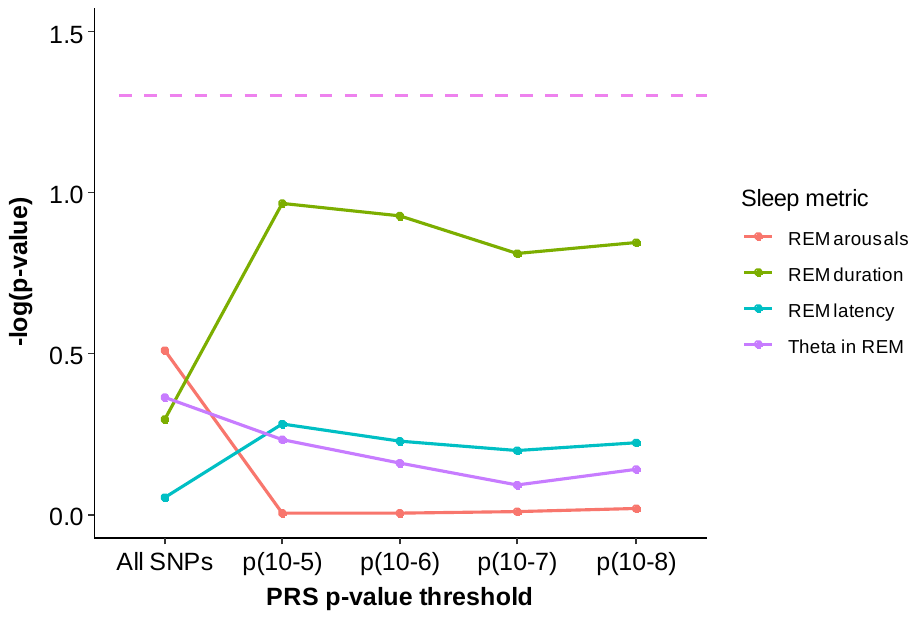

**Figure S3:** Associations between PRS for height and baseline night sleep metrics. Height was used as a negative control with the assumption of absence of any a priori association between the sleep phenotypes and a genetic liability for height. Plot shows statistical outcomes of GAMLSSs with four sleep metrics of interest versus PRS for height from conservative (p < 1 × 10^−8^) p value threshold to using all SNPs (N = 518). The models show absence of any significant association for PRS for height providing support to our findings. The models were corrected for sex, BMI, and TST or REM duration. The negative log transformation of p values of the associations is presented on the vertical axis. Horizontal line (pink dashed line) indicates p values threshold of 0.05.

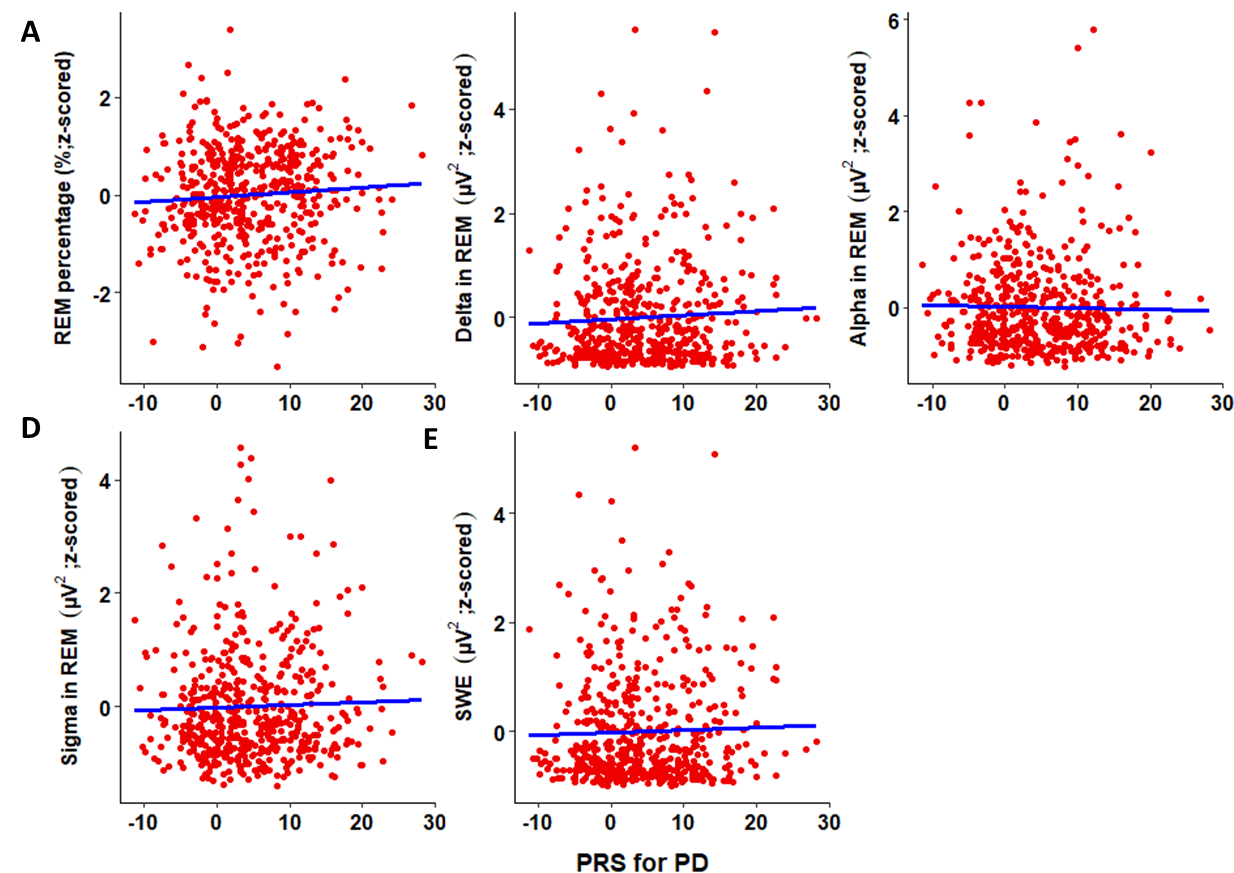

**Figure S4:** Associations between PRS for PD and other baseline night sleep metrics for specificity analysis. The associations between six additional sleep metrics during baseline night and PD PRS for SNPs at p-value threshold of 1x10^-8^ (*N* = 518). GAMLSS yielded significant associations only for REM percentage (**A**). Spearman correlation *r* is reported for completeness, refer to main text supplementary **Table S2** for statistical outputs of GAMLSSs. SWE: slow wave energy

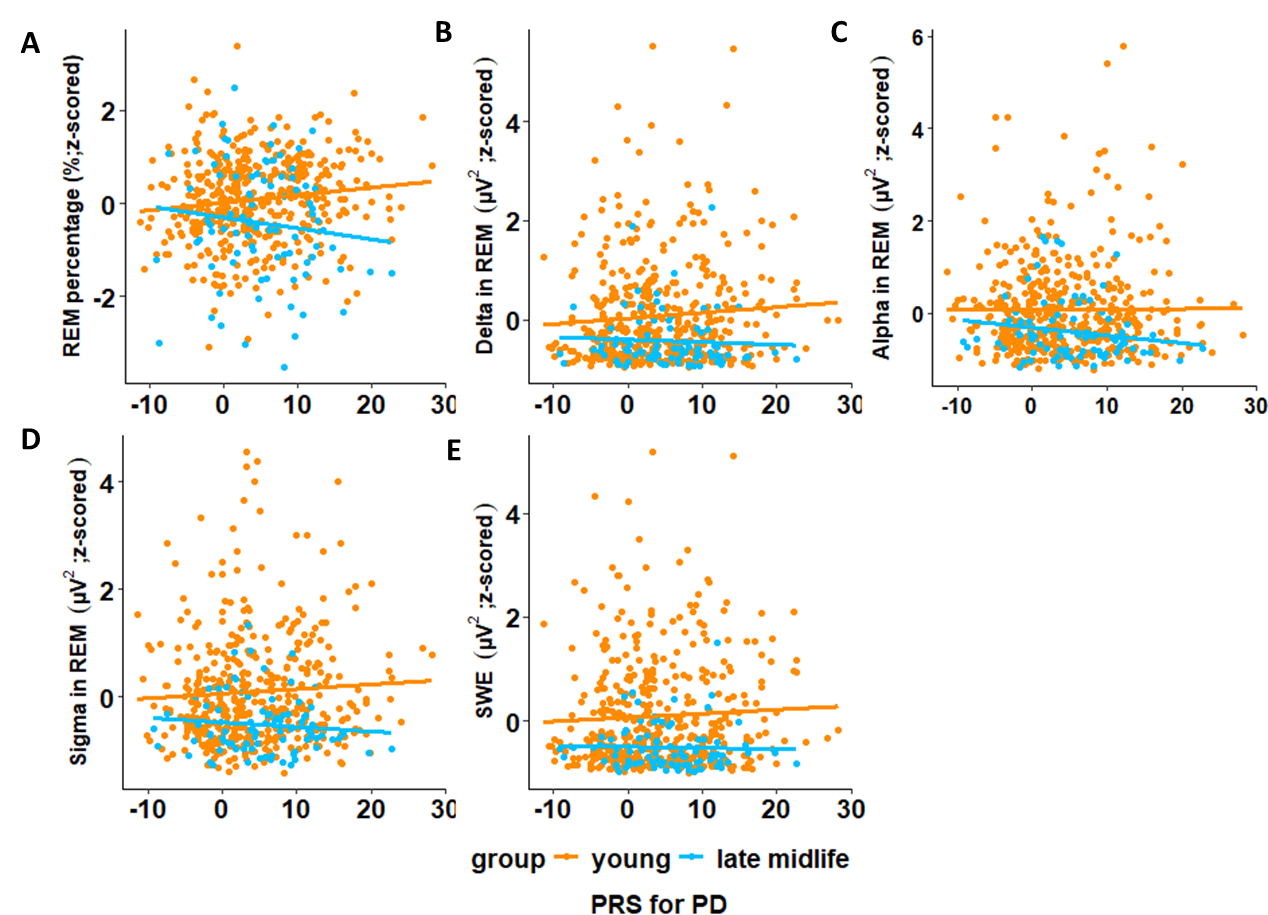

**Figure S5:** Age related associations between PRS for PD and other baseline night sleep metrics for young and late midlife sub-sample. **A.** The association between REM percentage during baseline night and PD PRS for SNPs at p-value threshold of 1x10^-8^ in younger sub-sample (*N* = 433) and late-midlife sub-sample (*N* = 85). GAMLSS yielded an interaction between PD PRS and age for REM sleep percentage in both young and late-midlife individuals (**p=0.003** and **p=0.010** respectively). These findings further support our previous results and imply that REM stage of sleep is associated with the PD risk with specific metrics playing a major role. Refer to main text **Table S3** for complete statistical outputs of GAMLSSs. **B-E.** GAMLSS with REM delta energy, REM alpha energy, REM sigma energy or slow wave energy (SWE) in NREM sleep showed no statistically significant association.

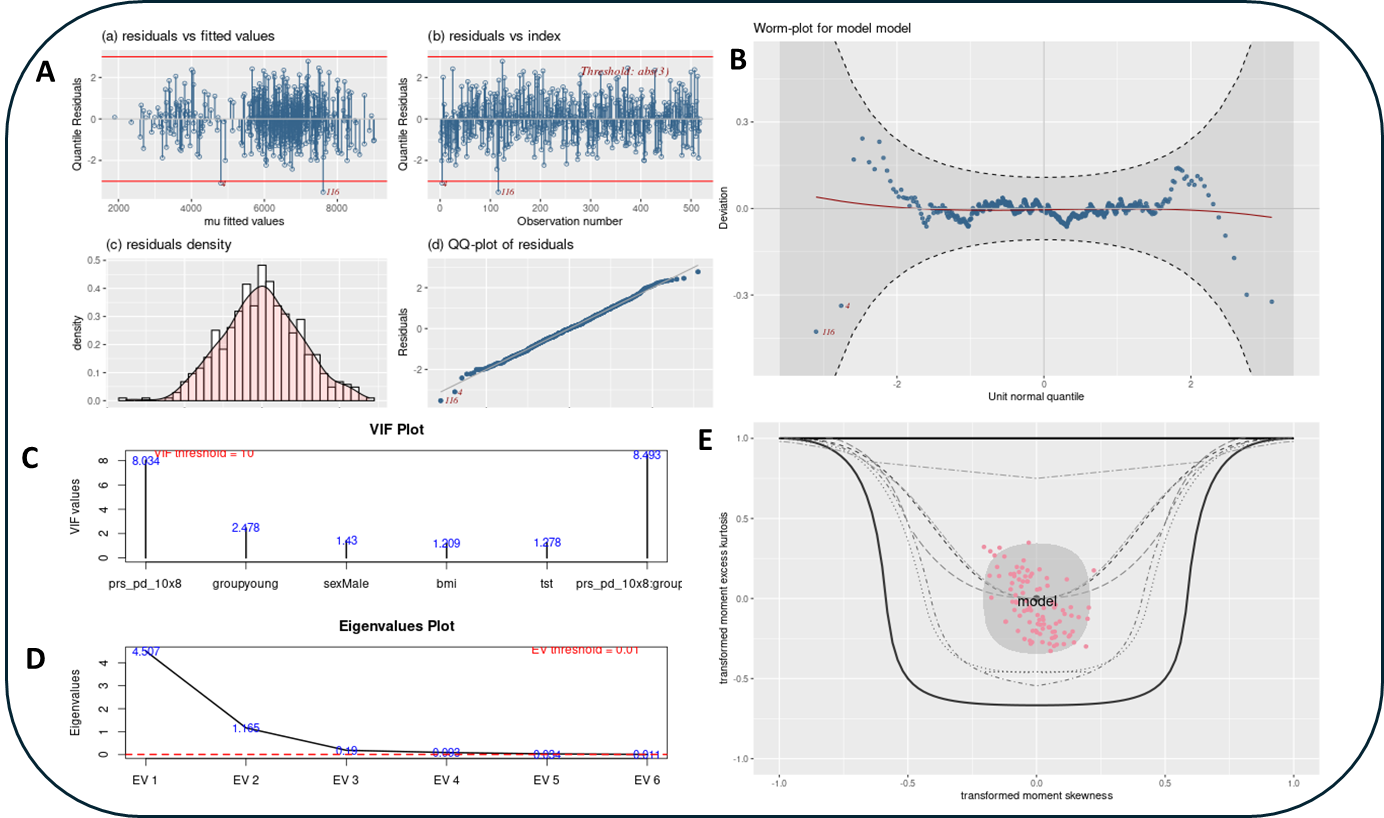

**Figure S6:** A representation of the GAMLSS model fitting diagnostics for REM theta as the dependent variable and PD PRS (10^-8^) as the main independent variable with age group as interactor and Sex, BMI, TST as covariates. **A.** (a) Residuals versus fitted values, and (b) residuals versus index, showing whether residuals appear randomly distributed. (c) Residual density and (d) QQ-plot of residuals, assessing normality. **B.** Worm plot, a refined QQ-plot highlighting deviations from normality. **C.** Variance inflation factor (VIF) plot and **D.** eigenvalues plot, checking for multicollinearity among predictors. **E.** Bucket (Moment skewness–kurtosis) plot, evaluating distributional shape. The figure shows that model fit was good with the selected family of distribution as identified by the fitdist function of GAMLSS. In the analysis, AIC, QQ plot and bucket plot (right lower panel was considered for selecting the best model.
